# Nuclear Mitochondrial Interaction Test Reveals Sex-Dependent Mitochondrial SNPs Interacting with Klotho Variants on Diabetes Risk

**DOI:** 10.64898/2026.02.12.705615

**Authors:** Tae Jung Oh, Hiroshi Kumagai, Kelvin Yen, Eileen M. Crimmins, Thalida E. Arpawong, Pinchas Cohen

## Abstract

**Context:** The environmental or other genetic factors might influence the effect of *Klotho* (*KL)* on glucose metabolism.

**Objective:** We investigated mitochondrial genetic variants that interact with *KL* single nucleotide polymorphisms (SNPs) to modulate diabetes risk.

**Methods:** We used the data from 7,047 non-Hispanic white participants of the Health and Retirement Study, a prospective observational study including adults aged 50 years and older from the United States. First, we performed single gene-wide association scans to identify *KL* SNPs associated with diabetes. Next, we performed a nuclear-by-mitochondrial interaction test (NuMIT) in which we use an identified *KL* SNP from the gene-wide scan to evaluate potential interactions with 85 mitochondrial SNPs in relation to diabetes.

**Results:** We failed to identify a significant association between diabetes and the *KL* SNP in our single gene-wide association test. However, we identified a novel variant (*KL* rs9563121) which showed a trend of increasing klotho mRNA levels with each additional minor allele. A NuMIT analysis identified mitochondrial SNPs, which showed significant interactions with rs9563121 in relation to diabetes risk. MitoG15929A showed significant interactions with rs9563121 in both men and women. MitoG15929A diminished the potential beneficial effect of *KL* rs9563121 on diabetes risk in women. Among men with the MitoG15929A variant, *KL* rs9563121 was associated with higher prevalence of diabetes.

**Conclusion:** The NuMIT approach revealed significant interactions between mitochondrial and nuclear DNA variants of *KL*. Furthermore, MitoG15929A may have a role in the interaction between diabetes and *KL* in a sex-dependent manner.

---

Mice with defects in the *Klotho* gene (*KL*) show effects on age-related phenotypes, such as short lifespans and arteriosclerosis (1), although *KL* overexpression extends lifespans (2). Therefore, *KL* has been thought to be an aging suppressor. Studies on the human genetic polymorphism of *KL* also suggest anti-aging effects of the Klotho protein in humans (3,4). A recent attempt to test the therapeutic potential showed that Klotho protein administration improved memory in old rhesus monkeys (5). In this context, genetic variations related to *KL* may not only represent a key pathological mechanism of human diseases in conjunction with aging but also have the potential to be therapeutic targets for human chronic diseases.

Type 2 diabetes can develop from imbalances between beta-cell dysfunction and insulin resistance, which are the core pathophysiologic defects of the disease (6). As aging is related to both beta-cell function and insulin resistance (7,8), type 2 diabetes is considered an age-related disease. However, the role of Klotho in glucose metabolism is not as simple as with other age-related phenomena such as longevity and cognition. *KL* mutant mice show decreases in insulin production but also increases in insulin sensitivity (9), which implicates action in the opposite direction for systemic glucose levels. Furthermore, overexpression of Klotho in mice pancreatic islets directly increased insulin production (10), while overexpression of *KL* in mice inhibits insulin signaling *in vivo* (2). Interestingly, diet can modify the influence of Klotho on glucose metabolism. For example, insulin sensitivity was lower in *KL* transgenic mice and higher in heterozygous mice compared to wild-type mice, respectively. Yet, under high-fat diets, insulin sensitivity was restored in *KL* transgenic mice and decreased in heterozygous mutant mice, a phenomenon that was reversed under the normal chow diet model (11). Therefore, environmental or other genetic factors, aside from *KL*, could influence the effect of Klotho on glucose metabolism. In this context, more precise evaluation is required to understand the role of Klotho on diabetes.

In addition, sex-dimorphism has been observed in the relationship between Klotho and glucose metabolism. For example, insulin resistance was observed exclusively in male transgenic mice and not in their female counterparts (2). Other human studies also showed sex differences in Klotho levels associated with certain clinical phenotypes of depression (12) and lipid profiles (13). This complex relationship between Klotho and human diseases prompted us to investigate a novel interaction involving *KL* genetic variations and diabetes considering potential interactions with sex. Furthermore, we included both nuclear and mitochondrial genomic information because most genetic studies analyzed the nuclear genome and mitochondrial genome separately, resulting in interactions between these two potentially being missed.

## RESEARCH DESIGN AND METHODS

### Study Population

The Health and Retirement Study (HRS) is a prospective observational study of community-dwelling adults over 50 years of age from the United States. The first recruitment was conducted in 1992, and the surveyed samples are more than 37,000 individuals. The study participants were followed up biannually. The cohort overview was published elsewhere (14). In the current study we used the data obtained in 2016. Race was self-reported and grouped as Hispanic, non-Hispanic Black, non-Hispanic White, and other. In this study, we analyzed data from non-Hispanic White individuals (n = 7,047) only because the number of subjects in other ethnic groups was relatively small (Non-Hispanic Black n = 1,686, Hispanic n = 1,288), but also due to the substantially different allelic frequencies across variants in the mitochondrial genome across ancestries. All participants provided written informed consent. The study was approved by the University of Southern California Institutional Review Board and the HRS cohort study was approved by University of Michigan Health Sciences Institutional Review Board.

### Definition of Diabetes, Study Covariates, and Genomic Data

Sociodemographic data were obtained by face-to-face interview or telephone. Diabetes was self-reported as diagnosed by a doctor or by the participants. Other variables included age, sex, body mass index, systolic and diastolic blood pressure, and comorbidities. Genotype data were accessed from the National Center for Biotechnology Information Genotypes and Phenotypes Database (dbGaP) (15). Genotyping was conducted on over 15,000 individuals using either the Illumina HumanOmni2.5-4v1 (2006 and 2008) and HumanOmni2.5-8v1 (2010) arrays and was performed by the NIH Center for Inherited Disease Research. Standard quality control procedures were implemented by the University of Washington Genetic Coordinating Center. Further detail is provided in the HRS documentation (16). Mitochondrial SNP alleles are binary coded as 0 or 1 whereas nuclear SNPs are coded as 0, 1 or 2 for the number of minor alleles.

### mRNA Expression Levels Using RNA-sequencing Data

In the HRS cohort, RNA samples were sequenced as 50 base pair single read sequences with a minimum of 20 million reads per sample on a NovaSeq with ribosomal and globin messenger RNA reduction completed prior to sequencing. Sequencing was based on the TopMed/GTEX analysis pipeline, first using the STAR aligner (17) to align RNAseq reads to the GrCh38 reference genome from GENCODE, then calculating quality control metrics using RNASeQC (18). SAMTools (19) and RSEM (20) were used to obtain gene read counts. Counts were then normalized using the size factor method implemented by the R package DESeq2 (21) to account for sequencing depth differences. Gene expression values were calculated as the log_2_ counts per million (log2cpm). We then extracted the expression value for the *KL* gene.

### Statistical Analysis

Descriptive analysis was performed by SPSS version 27.0 (IBM Co., Armonk, NY, USA) and genetic data were analyzed by PLINK. Continuous data are presented as mean and standard deviation or standard error of the mean, and categorical data are presented as counts with percentages. Differences between two groups were assessed using Student t-test or Chi-Square test. ANOVA test was applied for testing the differences of clinical characteristics and Klotho mRNA levels between three groups. Logistic regression analysis was used to examine the association between genetic variation in the *KL* gene and diabetes status, stratified by sex, because we observed a trend suggesting gene-sex interaction for diabetes. Each model adjusted for age, body mass index, prior stroke, prior angina, and eigenvectors 1-6. Benjamini-Hochberg (BH) False Discovery Rate (FDR) was calculated to adjust the *P* values for multiple comparisons. Covariates were chosen based on statistical significance and excluding multicollinearity tested by variance inflation factor. The Nuclear-by-Mitochondrial Gene Interaction Test (NuMIT) was conducted in PLINK 1.9 (22) with the nuclear SNP candidate used to test effect modification of each mitochondrial SNP on diabetes status, by adding a nuclearSNP*mitochondrialSNP interaction term to each model. Sex-specific models were run, adjusting for age, body mass index, prior stroke, prior angina, and eigenvectors 1-6. BH FDR was calculated for multiple test correction. Adjusted *P* values less than 0.05 were considered statistically significant.

## RESULTS

We analyzed 7,047 subjects and diabetes prevalence was 26.9% in men and 21.7% in women. Subjects with diabetes were older and had higher BMI, and diabetes related comorbidities (**Supplementary Table S1**).

### Association of *KL* SNPs with Diabetes Risk

First, we performed logistic regression analysis in each sex group (**Table 1**). There were no significant SNPs for diabetes after BH adjustment. There was no association between diabetes prevalence, *KL* rs9563121 variant, and Klotho mRNA in men (**Fig. 1A and 1B**). However, an inverse association was found in the context of the relationship between diabetes prevalence and the *KL* rs9563121 variant in women (**Fig. 1C)**. In addition, we found that the minor allele of *KL* rs9563121 was associated with a higher level of Klotho mRNA in women (**Fig. 1D**).

**Figure 1.**
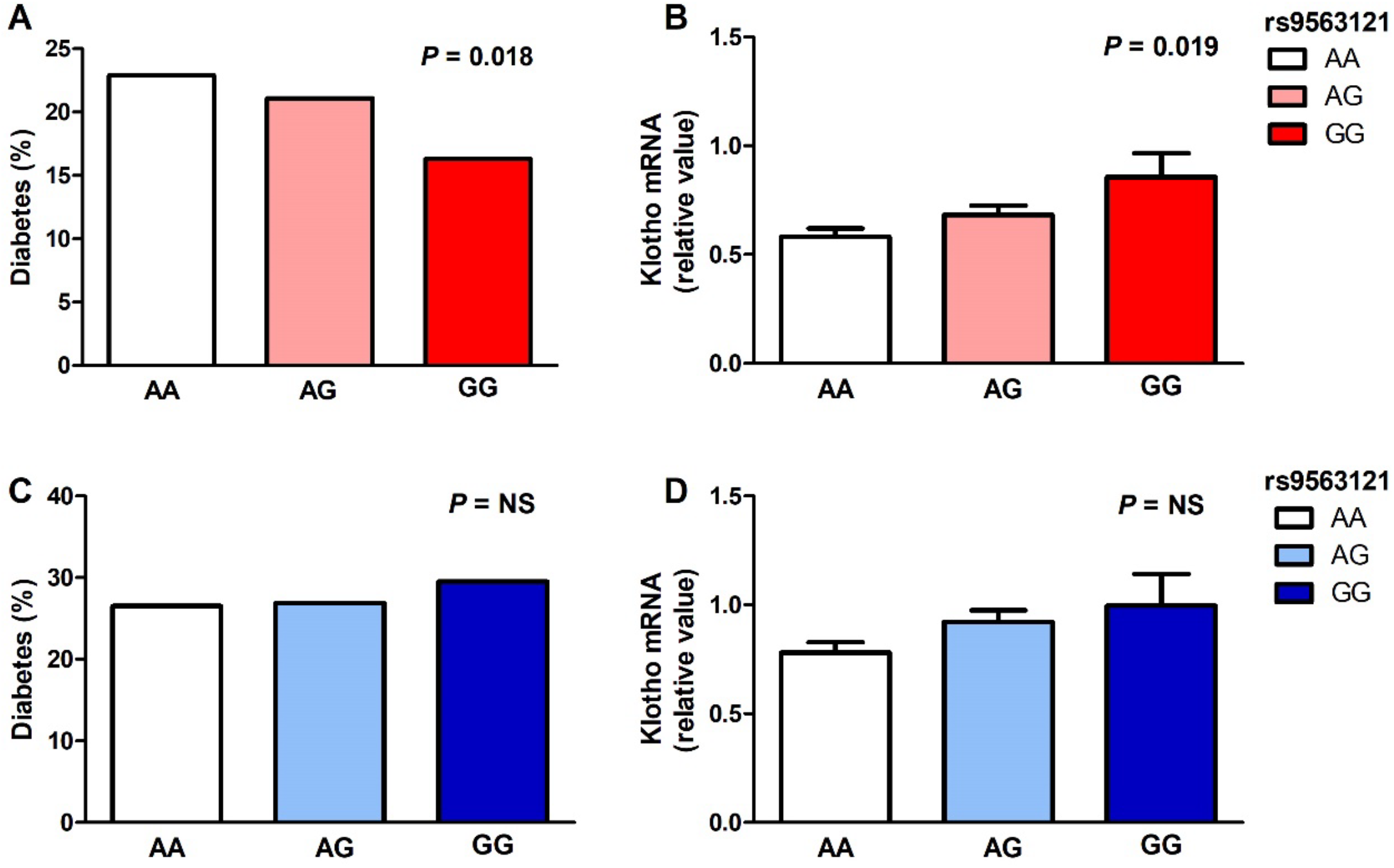
The prevalence of diabetes and Klotho mRNA levels by *KL* rs9563121 genotype. A. prevalence of diabetes and B. Klotho mRNA levels in women. C. prevalence of diabetes and D. Klotho mRNA levels in men.

### Nuclear Mitochondrial Gene Interaction Test (NuMIT) of *KL* SNP rs9563121

We tested whether the nuclear SNP of rs9563121 in the *KL* gene significantly interacted with mitochondrial SNPs to affect risk of diabetes (**Fig. 2**). This analysis termed NuMIT identified seven mitochondrial SNPs that significantly interacted with rs9563121 to affect diabetes risk. Among them, MitoG15929A significantly modified the association between *KL* rs9563121 and diabetes in males and females (**Fig. 3**). In men with a MitoG15929A minor allele, *KL* rs9563121 was associated with higher prevalence of diabetes (**Fig. 4A**). In contrast, the beneficial effect of the *KL* rs9563121 minor allele on diabetes was observed in women without MitoG15929A minor allele, but this effect diminished by MitoG15929A (**Fig. 4B**). However, MitoG15929A itself was not associated with diabetes in either men or women (data not shown).

**Figure 2.**
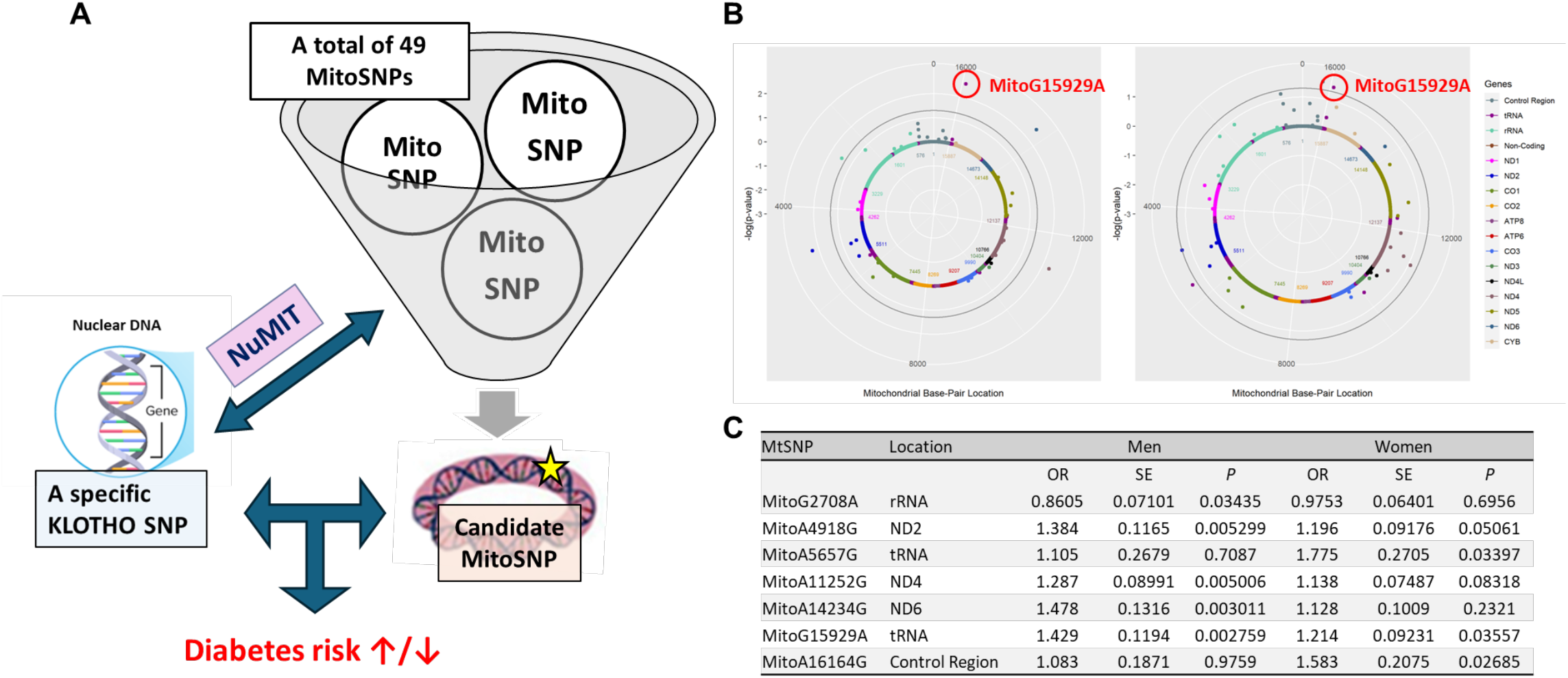
Nuclear by mitochondrial gene interaction test (NuMIT): An approach to identify mitochondrial variations that interact with *KL* variant rs9563121 to affect diabetes risk. A. Schematic figure of NuMIT. B. Solar plots presenting results of the nuclear by mitochondrial gene interaction test of *KL* rs9563121 with mitochondrial SNPs. Mitochondrial SNPs extending beyond the outer grey line are statistically significant by a permutation *P* value of 0.05. Red circles annotate a mitochondrial SNP with a *P* value less than 0.05 in both men and women. C. Summary of NuMIT between *KL* rs9563121 and mitochondrial SNPs. *P* value was adjusted by adjusted for age, body mass index, prior stroke, prior angina, and eigenvectors 1-6.

**Figure 3.**
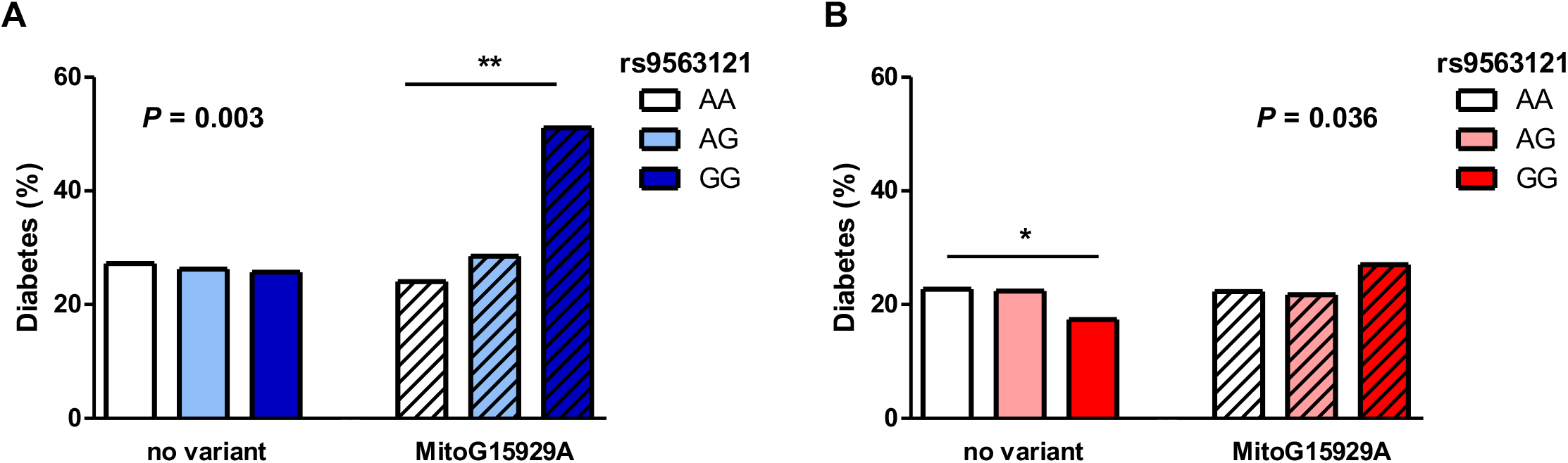
The effect of mitochondrial SNP G15929A (MitoG15929A) on diabetes risk depends on sex and *KL* SNP rs9563121. A Men. B Women. *P* values indicate the interaction between MitoG15929A and rs9563121 on diabetes risk. * *P*<0.05, ** *P*<0.01 for Chi-square test.

## CONCLUSIONS

Here, we report a novel interaction between a nuclear variant, *KL* SNP rs9563121, and mitochondrial SNP, MitoG15929A. NuMIT can be employed to discover novel genetic interactions within the context of each disease. The *KL* gene and diabetes are one example, and we successfully identified a novel mitochondrial variant through the NuMIT approach. Neither the *KL* SNP nor the Mito SNP showed a statistically significant association with diabetes alone. However, combining these two enabled us to identify an important modifier, MitoG15929A within the context of diabetes.

Previously genetic variations of *KL* have been reported to interact with another genetic polymorphism. For example, *KL* rs495392 has been reported to interact with *PNPLA3* rs738409, a well-known genetic determinant of fatty liver disease (23). In that study, the detrimental impact of *PNPLA3* rs738409 on severe hepatic steatosis was attenuated by *KL* rs495392. However, there was no report on nuclear and mitochondrial genetic interactions with *KL* polymorphisms. The genetic interaction between *KL* rs9563121 and MitoG15929A provides an example of how we can better understand nuclear and mitochondrial gene interactions that are associated with diabetes risk. Further studies are necessary to discover which peptides or molecules derived by this mitochondrial variant can interfere with the effect of Klotho. They might originate from mitochondria, known as mitochondrial-derived microproteins. Furthermore, it is crucial to further investigate the sex-specific effects of rs9563121 and MitoG15929A on glucose metabolism.

Klotho, as an aging suppressor, has been studied for its genetic variations in relation to various chronic diseases. Previous studies showed that rs9563121 was associated with posttraumatic stress disorder (24), while rs571118 and rs563925, which are in LD with rs9563121, showed delayed onset of end-stage renal disease (25) and low PSA levels (26), respectively. To the best of our knowledge, rs9563121 is reported here for the first time as a genetic variant related to diabetes. The *KL* rs9563121 was in high linkage disequilibrium with rs563925 and rs953124, both of which were suggestive of negative associations with diabetes in women based on raw p-values. This suggests these SNPs may be tagging another variant that was not genotyped on the array. A potential follow-up approach could involve using whole genome sequencing data to evaluate additional SNPs.

The *Klotho* gene encoded both transmembrane and circulating α-Klotho protein, and the circulating form can function as a hormone (27). In our study, we did not directly measure the circulating concentration of the Klotho protein. However, we can speculate about the level based on the mRNA expression levels in blood. Interestingly, the protective effect of Klotho was only observed in women. This might be why we observe differences in diabetes prevalence according to the presence of minor allele of *KL* rs9563121 in women. To further understand this sex-dimorphic effect of Klotho on diabetes, additional functional studies incorporating sex hormones will be necessary.

In our study, we have several limitations. First, given the nature of association study, we could not determine the causal effect of Klotho on diabetes. Second, our findings were driven by ethnic-specific phenomena. So, we cannot generalize our findings to other ethnic groups. Third, we did not validate our results using an independent cohort. Lastly, we proposed the interaction between nuclear and mitochondrial genes through statistical methods, but experimental research on actual biology would be necessary to validate this.

In conclusion, applying the novel NuMIT approach provided novel insight about the potential interaction between mitochondrial and nuclear DNA effects. MitoG15929A is the first identified mitochondrial variant to modify the risk of diabetes according to the *KL* rs9563121 variant and sex.

## Acknowledgments

We are sincerely grateful to the participants and researchers whose dedication to data collection and sharing made this study possible. We especially acknowledge the contributions of the Health and Retirement Study cohort members.

## Author Contributions

T.J.O., T.E.A., and P.C. conceived and designed the analysis. T.J.O, H.K., K.Y., E.M.C., T.E.A., and P.C. analyzed the data and interpreted the results. T.J.O, E.M.C., and T.E.A. provided statistical expertise. T.J.O., H.K., K.Y., T.E.A., and P.C. drafted the manuscript. All authors read and approved the final version of the manuscript.

## Abbreviations

BH: Benjamini-Hochberg
FDR: False Discovery Rate
HRS: Health and Retirement Study
KL: Klotho
MA: Minor allele
MAF: Minor allele frequency
NuMIT: Nuclear-by-Mitochondrial Gene Interaction Test

## References

1. Kuro-o M, Matsumura Y, Aizawa H, Kawaguchi H, Suga T, Utsugi T, Ohyama Y, Kurabayashi M, Kaname T, Kume E, Iwasaki H, Iida A, Shiraki-Iida T, Nishikawa S, Nagai R, Nabeshima YI. Mutation of the mouse klotho gene leads to a syndrome resembling ageing. Nature. 1997;390(6655):45–51.

2. Kurosu H, Yamamoto M, Clark JD, Pastor JV, Nandi A, Gurnani P, McGuinness OP, Chikuda H, Yamaguchi M, Kawaguchi H, Shimomura I, Takayama Y, Herz J, Kahn CR, Rosenblatt KP, Kuro-o M. Suppression of aging in mice by the hormone Klotho. Science. 2005;309(5742):1829–1833.

3. Zhu Z, Xia W, Cui Y, Zeng F, Li Y, Yang Z, Hequn C. Klotho gene polymorphisms are associated with healthy aging and longevity: Evidence from a meta-analysis. Mech Ageing Dev. 2019;178:33–40.

4. Mengel-From J, Soerensen M, Nygaard M, McGue M, Christensen K, Christiansen L. Genetic Variants in KLOTHO Associate With Cognitive Function in the Oldest Old Group. J Gerontol A Biol Sci Med Sci. 2016;71(9):1151–1159.

5. Castner SA, Gupta S, Wang D, Moreno AJ, Park C, Chen C, Poon Y, Groen A, Greenberg K, David N, Boone T, Baxter MG, Williams GV, Dubal DB. Longevity factor klotho enhances cognition in aged nonhuman primates. Nat Aging. 2023;3(8):931–937.

6. DeFronzo RA. Pathogenesis of type 2 diabetes mellitus. Med Clin North Am. 2004;88(4):787–835, ix.

7. Helman A, Avrahami D, Klochendler A, Glaser B, Kaestner KH, Ben-Porath I, Dor Y. Effects of ageing and senescence on pancreatic beta-cell function. Diabetes Obes Metab. 2016;18 Suppl 1:58–62.

8. Spinelli R, Baboota RK, Gogg S, Beguinot F, Bluher M, Nerstedt A, Smith U. Increased cell senescence in human metabolic disorders. J Clin Invest. 2023;133(12).

9. Utsugi T, Ohno T, Ohyama Y, Uchiyama T, Saito Y, Matsumura Y, Aizawa H, Itoh H, Kurabayashi M, Kawazu S, Tomono S, Oka Y, Suga T, Kuro-o M, Nabeshima Y, Nagai R. Decreased insulin production and increased insulin sensitivity in the klotho mutant mouse, a novel animal model for human aging. Metabolism. 2000;49(9):1118–1123.

10. Lin Y, Sun Z. Antiaging gene Klotho enhances glucose-induced insulin secretion by up-regulating plasma membrane levels of TRPV2 in MIN6 beta-cells. Endocrinology. 2012;153(7):3029–3039.

11. Gu H, Jiang W, You N, Huang X, Li Y, Peng X, Dong R, Wang Z, Zhu Y, Wu K, Li J, Zheng L. Soluble Klotho Improves Hepatic Glucose and Lipid Homeostasis in Type 2 Diabetes. Mol Ther Methods Clin Dev. 2020;18:811–823.

12. Zhang Y, Lu J, Huang S, Chen Y, Fang Q, Cao Y. Sex differences in the association between serum alpha-Klotho and depression in middle-aged and elderly individuals: A cross-sectional study from NHANES 2007-2016. J Affect Disord. 2023;337:186–194.

13. Lee J, Kim D, Lee HJ, Choi JY, Min JY, Min KB. Association between serum klotho levels and cardiovascular disease risk factors in older adults. BMC Cardiovasc Disord. 2022;22(1):442.

14. Sonnega A, Faul JD, Ofstedal MB, Langa KM, Phillips JW, Weir DR. Cohort Profile: the Health and Retirement Study (HRS). Int J Epidemiol. 2014;43(2):576–585.

15. dbGaP, Health and Retirement Study. National Center for Biotechnology Information: Bethesda, MD. 2012.

16. HRS, Quality control report for genotypic data. University of Washington. St. Louis, MO. 2012:44.

17. Dobin A, Davis CA, Schlesinger F, Drenkow J, Zaleski C, Jha S, Batut P, Chaisson M, Gingeras TR. STAR: ultrafast universal RNA-seq aligner. Bioinformatics. 2013;29(1):15–21.

18. DeLuca DS, Levin JZ, Sivachenko A, Fennell T, Nazaire MD, Williams C, Reich M, Winckler W, Getz G. RNA-SeQC: RNA-seq metrics for quality control and process optimization. Bioinformatics. 2012;28(11):1530–1532.

19. Li H, Handsaker B, Wysoker A, Fennell T, Ruan J, Homer N, Marth G, Abecasis G, Durbin R, Genome Project Data Processing S. The Sequence Alignment/Map format and SAMtools. Bioinformatics. 2009;25(16):2078–2079.

20. Li B, Dewey CN. RSEM: accurate transcript quantification from RNA-Seq data with or without a reference genome. BMC Bioinformatics. 2011;12:323.

21. Love MI, Huber W, Anders S. Moderated estimation of fold change and dispersion for RNA-seq data with DESeq2. Genome Biol. 2014;15(12):550.

22. Chang CC, Chow CC, Tellier LC, Vattikuti S, Purcell SM, Lee JJ. Second-generation PLINK: rising to the challenge of larger and richer datasets. Gigascience. 2015;4:7.

23. Liu WY, Zhang X, Li G, Tang LJ, Zhu PW, Rios RS, Zheng KI, Ma HL, Wang XD, Pan Q, de Knegt RJ, Valenti L, Ghanbari M, Zheng MH. Protective association of Klotho rs495392 gene polymorphism against hepatic steatosis in non-alcoholic fatty liver disease patients. Clin Mol Hepatol. 2022;28(2):183–195.

24. Wolf EJ, Morrison FG, Sullivan DR, Logue MW, Guetta RE, Stone A, Schichman SA, McGlinchey RE, Milberg WP, Miller MW. The goddess who spins the thread of life: Klotho, psychiatric stress, and accelerated aging. Brain Behav Immun. 2019;80:193–203.

25. Bostrom MA, Hicks PJ, Lu L, Langefeld CD, Freedman BI, Bowden DW. Association of polymorphisms in the klotho gene with severity of non-diabetic ESRD in African Americans. Nephrol Dial Transplant. 2010;25(10):3348–3355.

26. Kim HJ, Lee J, Lee SY, Cheong HS, Kye YS, Kim W, Byun SS, Myung SC. The association between KL polymorphism and prostate cancer risk in Korean patients. Mol Biol Rep. 2014;41(11):7595–7606.

27. Landry T, Shookster D, Huang H. Circulating alpha-klotho regulates metabolism via distinct central and peripheral mechanisms. Metabolism. 2021;121:154819.

